# The myotendinous junction marker collagen XXII enables zebrafish postural control learning and optimal swimming performance through its force transmission activity

**DOI:** 10.1101/2021.07.14.452354

**Authors:** Marilyne Malbouyres, Alexandre Guiraud, Christel Lefrançois, Mélanie Salamito, Pauline Nauroy, Laure Bernard, Frédéric Sohm, Bruno Allard, Florence Ruggiero

## Abstract

Although the myotendinous junction (MTJ) is essential for skeletal muscle integrity, its contribution to skeletal muscle function remains largely unknown. Here, we show that CRISPR-Cas9-mediated gene ablation of the MTJ marker *col22a1* in zebrafish identifies two distinctive phenotypic classes: class 1 individuals reach adulthood with no overt muscle phenotype while class 2 display severe movement impairment and eventually dye before metamorphosis. Yet mutants that are unequally affected are all found to display defective force transmission attributed to a loss of ultrastructural integrity of the MTJ and myosepta, though with distinct degrees of severity. The behavior-related consequences of the resulting muscle weakness similarly reveal variable phenotypic expressivity. Movement impairment at the critical stage of swimming postural learning eventually causes class 2 larval death by compromising food intake while intensive exercise is required to uncover a decline in muscle performance in class 1 adults. By confronting MTJ gene expression compensation and structural, functional and behavioral insights of MTJ dysfunction, our work unravels variable expressivity of *col22a1* mutant phenotype. This study also underscores *COL22A1* as a candidate gene for myopathies associated with dysfunctional force transmission and anticipates a phenotypically heterogeneous disease.

## Introduction

The myotendinous junction (MTJ) is a specialized anatomical region where tendon collagen fibers insert into the muscle basement membrane. These muscle-tendon connection sites are appropriately located and organized to transmit muscle contractile forces to tendons and create a movement. This is mainly due to the presence of two independent trans-sarcolemmal linkage systems that structurally link the intracellular contractile elements of muscle cells to the extracellular matrix (ECM) of tendons, the dystrophin-glycoprotein complex (DGC) and the α7β1 integrin complex. Mutations in their associated genes, such as dystrophin, α-7integrin, and α-2 laminin have been associated with myopathies in patients and animal models revealing that most of the MTJ components are unconditionally required for proper muscle function and integrity (Bassett et al, 2003; Postel et al, 2008; Hall et al., 2007). The impact of these mutations in MTJ formation and/or function remains poorly documented, particularly in the context of human diseases for which biopsies at this particular location would be too prejudicial for the patients. Yet mice that lack dystrophin (mdx; Dmd) (Law and Tidball, 1993; Law et al., 1995), laminin α2 (dy; Lama2) (Desaki, 1992) or integrin α7 (Miosge et al., 1999; Welser et al., 2009) all exhibit a striking reduction in the number of membrane folds at MTJ. Zebrafish has proven instrumental in the study of MTJ components in developing skeletal muscle. In zebrafish, mutant lines of the structural proteins enriched at the MTJ (*sapje/dystrophin, caf/lama2* and *patchytail/dag1*) or morpholino-knockdown of *itg7a* and *col22a1* in zebrafish embryos all displayed compromised muscle attachments that result in muscular dystrophies of various severities (Ingham, 2009; Sztal et al., 2012; Charvet et al, 2013). However *in vivo* function of these components was generally limited to zebrafish larval stage while muscular dystrophies are often progressive muscle disorders.

Collagen XXII is a recognized marker of the MTJ that was first described by Koch and colleagues (Koch et al, 2004). Functional analysis in zebrafish showed that *col22a1* expression concentrates at the ends of muscle fibers guiding the protein deposition at the junctional ECM. ECM proteome across muscle-tendon interface found collagen XXII restricted to the muscletendon junction tissue (Jacobson et al, 2020). Single-nucleus RNA-seq analysis in mice recently suggested that myonuclei near the muscle ends and tenocytes may both contribute this collagen (Petrany et al, 2020). While ColXXII binding partners have not been biochemically identified yet, synergistic interactions suggested that collagen XXII is a constitutive protein of the transmembrane α7β1 linkage system (Charvet et al, 2013) and electron microscopy revealed that ColXXII is located at the outer surface of the MTJ suggesting a structural role linking the basement membrane to the tendinous ECM (Koch et al, 2004; Charvet et al, 2013). As such, ColXXII could be the missing link that anchors muscle basement membrane network to the tendon collagen fibers. Although recognized as an important structural protein of the MTJ as recently exemplified in humans where high mRNA expression of *COL22A1* at the MTJ is associated with muscle injury risk in athletes (Miyamoto-Mikami et al, 2020), the role of this ECM protein in adult muscle function and performance remains overlooked.

Here we generated loss-of-function mutants in zebrafish *col22a1* using CRISPR/Cas9 technology to detail the function of ColXXII beyond early larval stages. Phenotype discrepancy between morphants and knock-out stable lines have been previously reported (Schulte-Mecker and Stainier, 2014; Kok et al, 2015; El-Brolosy and Stainier, 2017). But, as a rare example of a strong correlation between morpholino-induced and mutant phenotypes (Schulte-Mecker and Stainier, 2014), we further documented the requirement of collagen XXII for proper development of the musculotendinous system, for contractile force transmission and ultimately movements not only in larvae but in adults. By combining ultrastructural analyses, muscle performance measurements and behavioral assays, we demonstrate that the lack of ColXXII results in the systematic dysfunction of musculotendinous system that impairs postural behavior learning and swimming performance but with clear penetrance variability as often observed in human disease.

## Results

### Ablation of the MTJ marker *col22a1* results in two distinct phenotypic classes in the two independent mutant lines

ColXXII morphant embryos showed dystrophic-like phenotype (Charvet et al, 2013). But muscular dystrophy demonstrates marked skeletal muscle phenotypic modulation (Chelly and Desguerre, 2013). Thus, to further analyze the function of *col22a1* in larvae and adults, we generated two distinct *col22a1* knockout lines using CRISPR/Cas9 technology: *col22a1^vWA^ and col22a1^TSPN^* lines that target exon 2 and exon 6, respectively (Figure 1A - Figure supplement 1). Whole-mount immunostaining with antibodies against zebrafish ColXXII (Charvet et al, 2013) confirmed the complete extinction of ColXXII expression in homozygous *col22a1^vWA−/−^* and *col22a1^TSPN−/−^* mutants (Figure 1B).

**Figure 1:**
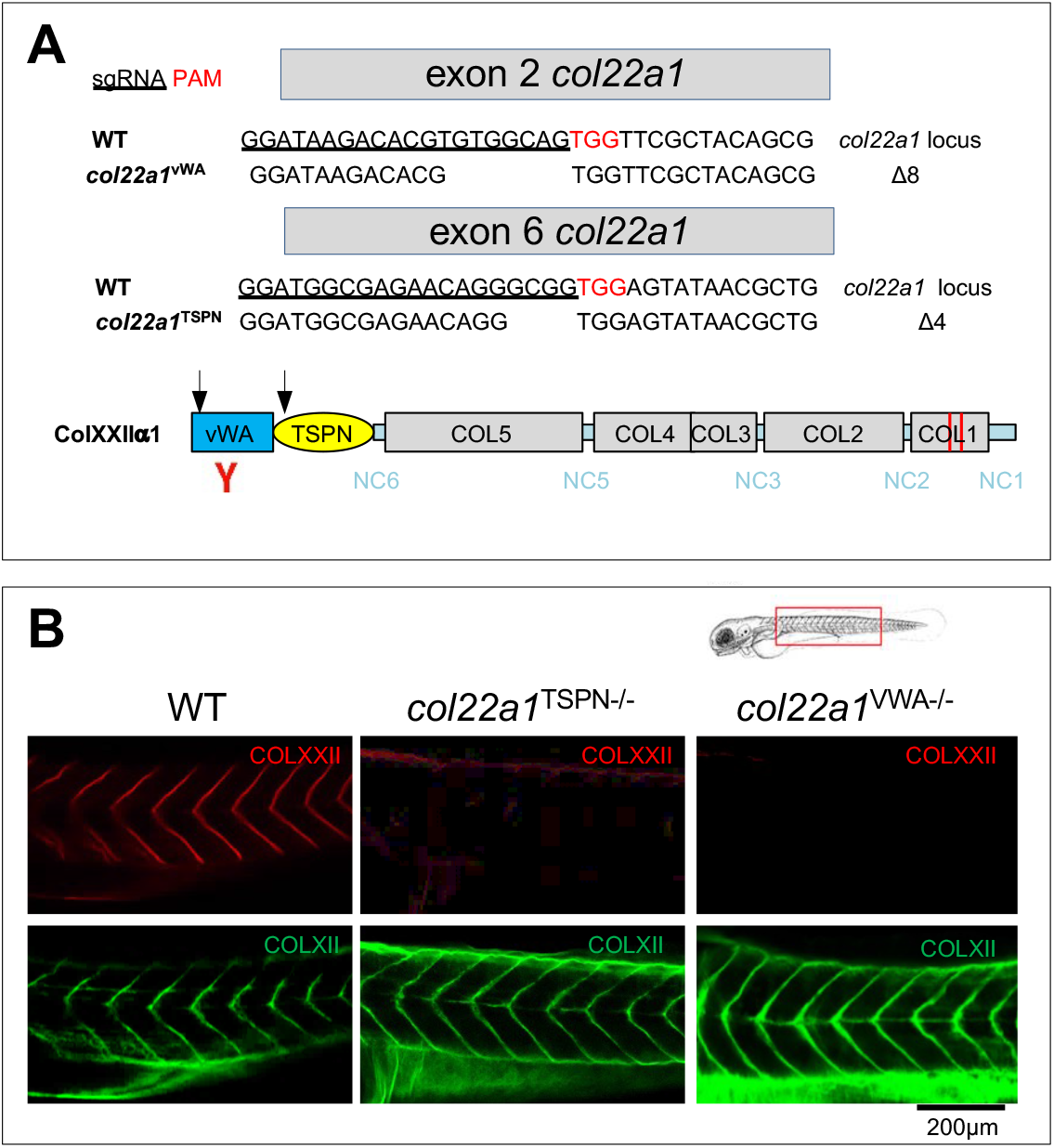
Generation and validation of *col22a1 knockout zebrafish lines*. A. Upper panel. Generation of the CRISPR/Cas9-mediated *col22a1* knockout zebrafish lines, *col22a1^vWA^* and *col22a1^TSPN^*. Schematic representations of the targeted *col22a1* regions in *col22a1* exon 2 (*col22a1^vWA^* line) and exon 6 (*col22a1^TSPN^* line). For each exon, the upper sequence corresponds to the WT sequence containing the sgRNA-targeted nucleotides (underlined) followed by the PAM sequence (in red). The lower sequence corresponds to the resulting 8 nucleotides and 4 nucleotides deletions in exon 2 (*col22a1^vWA^* line) and exon 6 (*col22a1^TSPN^* line), respectively. Each deleted nucleotide is represented by a space. Lower panel shows the zebrafish collagen XXII (ColXXII) primary structure. At the N-terminus, the VWA and TSPN domains represent the building blocks of the NC6 domain. COL1-COL5, collagenous domains, NC1-6, non-collagenous domains. Red bars are short interruptions in COL1. In contrast to humans, zebrafish *col22a1* lacks a NC4 domain (resulting in a fused COL3-COL4 domain). Y (in red) points to the recombinant domain used to generate antibodies to ColXXII (Charvet el al, 2013). Black arrows indicate the predicted truncation of ColXXII resulting from premature codon stop in exon 2 (*col22a1^vWA^* line) and exon 6 (*col22a1^TSPN^* line) domains. B. Whole-mount immunofluorescence staining of 5 dpf WT (left), *col22a1*^*TSPN*−/−^ (middle) and *col22a1*^*vWA*−/−^ (right) with anti-ColXXII (red) and the myoseptal marker anti-ColXII (green) as positive control. The upper cartoon shows orientation of the 5pdf larvae, the red box indicates the region imaged in WT, *col22a1*^*TSPN*−/−^ and *col22a1*^*vWA*−/−^ larvae. dpf: days post-fertilization.

At first, the *col22a1^vWA−/−^* and *col22a1^TSPN−/−^* fish appeared to develop normally and did not show any obvious muscular phenotype during embryo development. They hatched normally and successfully reached adulthood with no visible defects to the naked eye. If not genotyped, homozygous and heterozygous fish at 5 dpf were phenotypically indistinguishable from their WT siblings. However, a closer examination of the survival rate of the mutant fish astonishingly revealed that about 30% of the fish between 5dpf and adult stage were lost in clear contrast with the average mortality observed in WT animals (2-3%) (Table 1). This percentage did not vary when *col22a1^TSPN−/−^* or *col22a1^vWA−/−^* individuals that have reached adulthood were incrossed (Table 1). A closer examination of larvae during their growth revealed that, while most mutant fish remained phenotypically normal and reach adulthood, a subset of the total population started to show discernable musculoskeletal abnormalities at approximately 2 weeks post-fertilization (wpf) (Figure 2A). Because this variability could be due to a difference in the RNA decay efficiency that might not have cleaned of all defective transcripts, we measured *col22a1* transcript levels on RNA extracts of trunks of the two phenotypic subpopulations of larvae with q-PCR. We showed that *col22a1* transcript levels dropped to almost zero in the two phenotypic subpopulations of the mutant lines (Figure 2C). The severe phenotypic traits discernable at about 2 wpf larvae included impaired trunk movement and swimming capacities (Videos 1-3) and a slight but significant reduction of the total body length (Figure 2A and B, quantification). These larvae prematurely died shortly afterward (Figure 2A and Table 1). The cause of their death, few hours after the critical 2wpf stage is not known. One likely explanation is the compromised food intake due to their impaired motility as we observed that the intestines of these larvae were optically empty compared to WT and homozygous mutants with no overt phenotype (Figure 2A).

**Table 1:** Variable expressivity of *col22a1^vWA−/−^* and *col22a1^TSPN−/−^* larval phenotypes at 2wpf. The severity of the phenotype is scored as swimming capacity and posture impairment using a stereoscope. dpf, days post-fertilization. Numbers into brackets indicate that different laying from F5 fish were used.

| Genotype | Generation | Number of embryos | Larval phenotype (%) |  |  | Premature mortality rate (%)<br>(before 4dpf) |
| --- | --- | --- | --- | --- | --- | --- |
|  |  |  | Wildtype | Class 1 | Class 2 |  |
| WT | F4 | 214 | 96.73 | - | - | 3.27 |
| WT | F5 | 225 | 98.22 | - | - | 1.78 |
| WT | F6 | 343 | 97.08 | - | - | 2.92 |
| *col22a1*<sup>vWA-/-</sup> | F5 (1) | 88 | - | 67.05 | 31.82 | 1.13 |
| *col22a1*<sup>vWA-/-</sup> | F5 (2) | 113 | - | 74.34 | 23.89 | 1.77 |
| *col22a1*<sup>vWA-/-</sup> | F6 | 84 | - | 67.86 | 29.76 | 2.38 |
| *col22a1*<sup>TSPN-/-</sup> | F5 (1) | 109 | - | 68.81 | 29.36 | 1.83 |
| *col22a1*<sup>TSPN-/-</sup> | F5 (2) | 141 | - | 72.34 | 25.53 | 2.13 |
| *col22a1*<sup>TSPN-/-</sup> | F6 | 90 | - | 63.33 | 35.56 | 1.11 |

**Figure 2:**
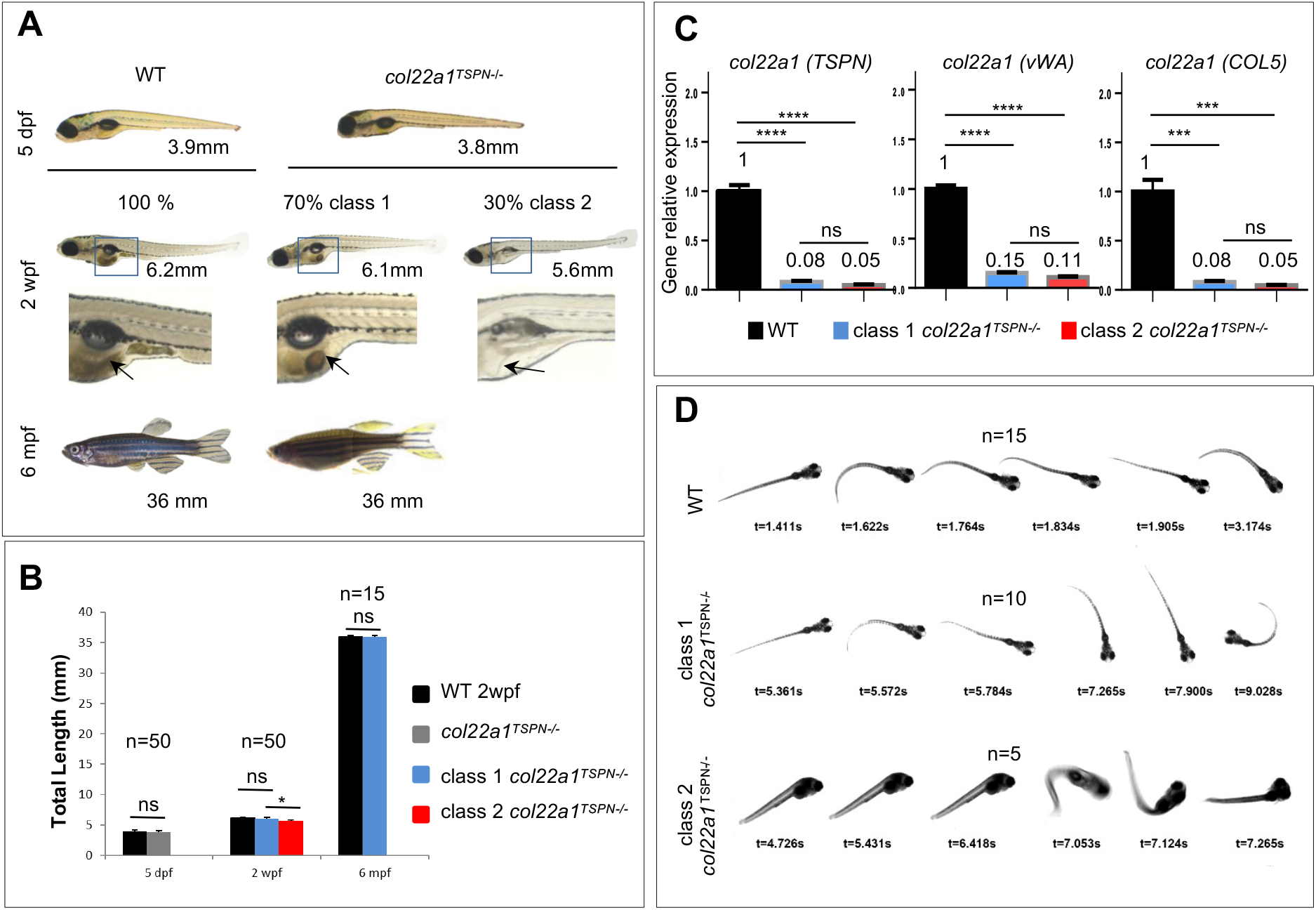
Knockout of *col22a1* results in class 1 and class 2 *phenotypic classes*. A. Representative bright field images of 5 dpf, 2 wpf and 6 mpf *col22a1*^*TSPN*−/−^ and WT individuals. *col22a1*^−/−^ class 2 larvae can be identified at around 2 wpf based on their posture defect and altered swimming capacity compared to WT. Arrows point to the empty gut observed only in 2 wpf *col22a1*^*TSPN*−/−^ class 2 fish. Values (in mm) are the total body length of individual fish. B. Quantification of the total body length of 5dpf *col22a1^TSPN−/−^* larvae (grey bars), 2 wpf and 6mpf class 1 (blue bars) and class 2 (red bars) *col22a1^TSPN−/−^* fish and WT (black bars). C. Real time RT-PCR analysis of *col22a1* gene expression in 2 wpf class 1 (blue bars) and class 2 (red bars) *col22a1^TSPN−/−^* fish and WT (black bars). Primers have been designed in different *col22a1* gene as indicated into brackets (TSPN, vWA and COL5 domains). Data were normalized to *polr2d*. Levels of *col22a1* expression were compared to WT expression, which was arbitrary set to 1. Data are mean ±SEM (n=3); ****p<0.0001; ***p<0,001; **p<0,01; *p<0,05; ns, not significant. D. Postural and swimming behaviors of 2wpf larvae. Single frame from videos of 2 wpf *col22a1*^*TSPN*−/−^ class 1 (row 2) and class 2 (row 3) larvae and WT (row 1). t, time expressed in seconds (s). n= number of analyzed larvae; dpf, days post-fertilization; wpf, weeks post-fertilization; mpf, months post-fertilization.

In conclusion, lack of *col22a* in zebrafish generates two distinct phenotypic classes in the progeny at the extremes of the potentially wide musculoskeletal phenotype spectrum: “class 1”, showing no overt phenotype and “class 2” that prematurely dies at the critical premetamorphosis stage. This phenotypic variability of the *col22a1* mutants can result from variable expressivity or incomplete penetrance.

### Movement dysfunction in class 2 larvae results in ineffective postural swimming bouts critical for larval survival

We next decided to investigate the cause of the class 2 larva death before metamorphosis. Interestingly, observations of freely swimming larvae and movies strikingly revealed that a subpopulation of larvae from both *col22a1^TSPN−/−^* and *col22a1^vWA−/−^* larvae failed to maintain horizontal posture and displayed defects in buoyancy at a stage they should have acquired this behavioral capacity crucial for their survival (Gla07sauer and Straka, 2017; Ehrlich and Schoppik, 2017) as in WT siblings (Videos 1-3). Instead, class 2 larva stay in a vertical position because they are unable to compensate the downward drag of the head by initiating bouts, likely because of defective muscle contractile response (Video 3 compared to Video 1). In addition, when some affected individuals managed to swim, they never succeeded to reach out the microscope field of view, in striking contrast to 2wpf WT or mutant individuals with no overt motility phenotype (class 1) that reached out the observation field of view several times in the same time window (Videos 1-3). Images extracted from videos detailed the erratic movements and the postural defects of the *col22a1^TSPN−/−^* and *col22a1^vWA−/−^* mutants compared to WT and class 1 mutant siblings (Figure 2D- Figure supplement 2). As *col22a1^TSPN−/−^* and *col22a1^vWA−/−^* mutants result in the same phenotypic classes in the same proportion, when the two lines were indifferently used, they are hereafter named *col22a1^−/−^* to facilitate the reading.

In conclusion, movement problems that seriously affect “class 2” larvae prevent them to acquire horizontal swimming posture that may correlate with their inability to respond to swimming bouts at a critical time for larval survival. This confirms the cause of their death as the reduced swimming capacity of the larvae critically compromises food-intake, explaining the abnormal high frequency of empty-gut class 2 larvae (Figure 2A).

### *col22a1^−/−^* mutants show similar musculoskeletal phenotype but with variable expressivity

We next hypothesize that the striking difference in the severity of the phenotypes could be the result of incomplete penetrance or variable expressivity of a musculoskeletal phenotype. The class 2 severe phenotype may correlate with the inability of the mutant musculoskeletal muscle system to appropriately respond to swimming bouts elicited by the larvae to maintain their horizontal position and the apparent absence of phenotype in class 1 fish may reflect a mild musculoskeletal phenotype that cannot be overtly seen. To address this question, we decided to measure electrically evoked muscle force generation of *col22a1^TSPN−/−^* mutant fish that grow to adulthood (class 1) and compare the values to *col22a1^TSPN-/+^* and WT siblings and the subset of *col22a1^TSPN−/−^* larvae that display severe musculoskeletal defects at 2 wpf (class 2). Interestingly, the contractile force generated by *col22a1^TSPN−/−^* muscles from both phenotypic classes was considerably lower compared to WT siblings and the mean amplitude of the twitch response was about two to four times smaller than WT in *col22a1^TSPN−/−^* phenotypic classes, class 1 and class 2, respectively (Figure 3A). Tetanic forces were also significantly lower in class 1 *col22a1^TSPN−/−^* larvae with no overt musculoskeletal phenotype compared to WT and heterozygote fish, but again, class 2 larvae with early motility-defective phenotype showed even lower values (Figure 3B). In that, our findings were reminiscent to variable expressivity and not incomplete penetrance. We conclude that lack of ColXXII impacts the efficacy of muscle contraction and force in all larvae but with variable expressivity and not because of incomplete penetrance.

**Figure 3:**
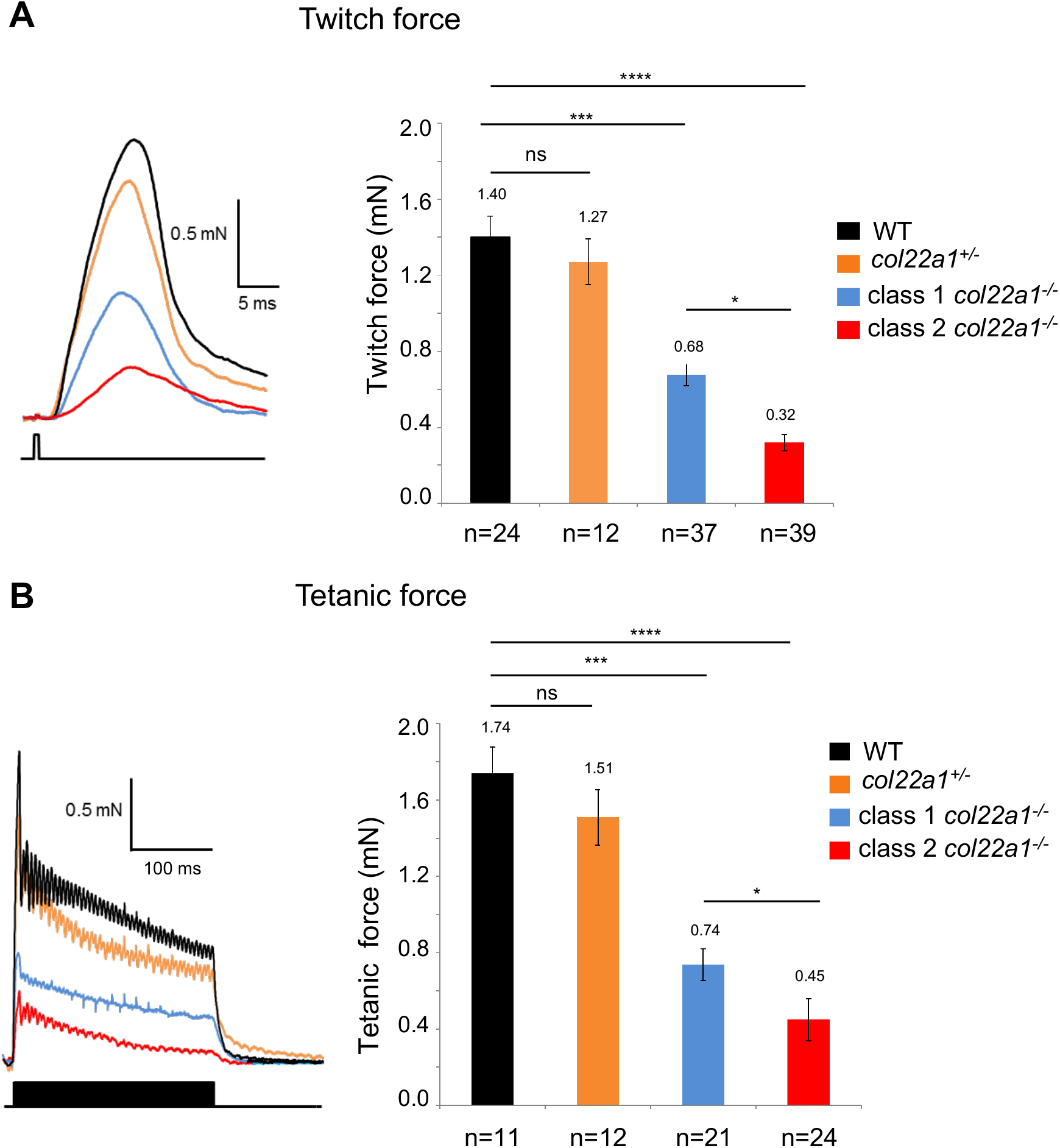
Muscle contractile performance of mutant larvae at 2wpf. A. Representative twitch responses (upper traces) elicited by single supramaximal electric shocks of 0.5 ms duration (lower trace) in *col22a1*^*TSPN*−/−^ class 1 (blue), class 2 (red), *col22a1*^*TSPN*+/-^ (orange) and wild-type (black) larvae (left panel) and histograms showing mean twitch amplitudes in different fish lines (right panel). B. Representative tetanic force (upper traces) recorded in response to trains of 0.5 ms duration electric shocks delivered at a frequency of 50 Hz during 250 ms (lower trace) (left panel) and histograms showing mean amplitude of tetanic force in different fish lines (right panel). Data are means ± SEM.****p<0.0001; ***p<0,001; **p<0,01; *p<0,05; ns, not significant. n = number of analyzed larvae.

We previously postulated that the observed reduced muscle performance of 5 dpf *col22a1* morphants is the consequence of distended vertical myosepta and reduced MTJ folds that cannot transmit properly the muscle contractile force (Charvet et al, 2013). Thus, we next analyzed the ultrastructure of the junctional region between skeletal muscles from mutants showing severe or mild phenotype at 2wpf. Interestingly, 2wpf *col22a1^TSPN−/−^* trunk muscles revealed structural defects in the MTJ with a range of severity between class 1 and 2 phenotypes. The differentiation and overall organization of sarcomeres as observed with TEM appeared not affected by the lack of collagen XXII in both phenotypic classes of *col22a1^TSPN−/−^* mutants (Figure 4B and C compared to A). However, all 2 wpf mutants showed a reduced number of the characteristic MTJ finger-like folds and distended myosepta that contained loosely-organized collagen fibrils (class1; Figure 4E and class 2; Figure 4F) compared to controls (Figure 4D) but these defects are much more severe in class 2 than in class 1 mutants. More specific to the class 2 mutants was the lack of muscle cohesiveness next to the MTJ (Figure 4F compared to D,E), but in striking contrast to *col22a1* morphants (Charvet et al, 2013), only rare muscle fiber detachment at the MTJ site was observed with histology (not shown). Altogether, our data show here that lack of ColXXII at the MTJ results in muscle force weakness due to compromised contractile force transmission of contractile strength from muscle to myoseptal tendon.

**Figure 4:**
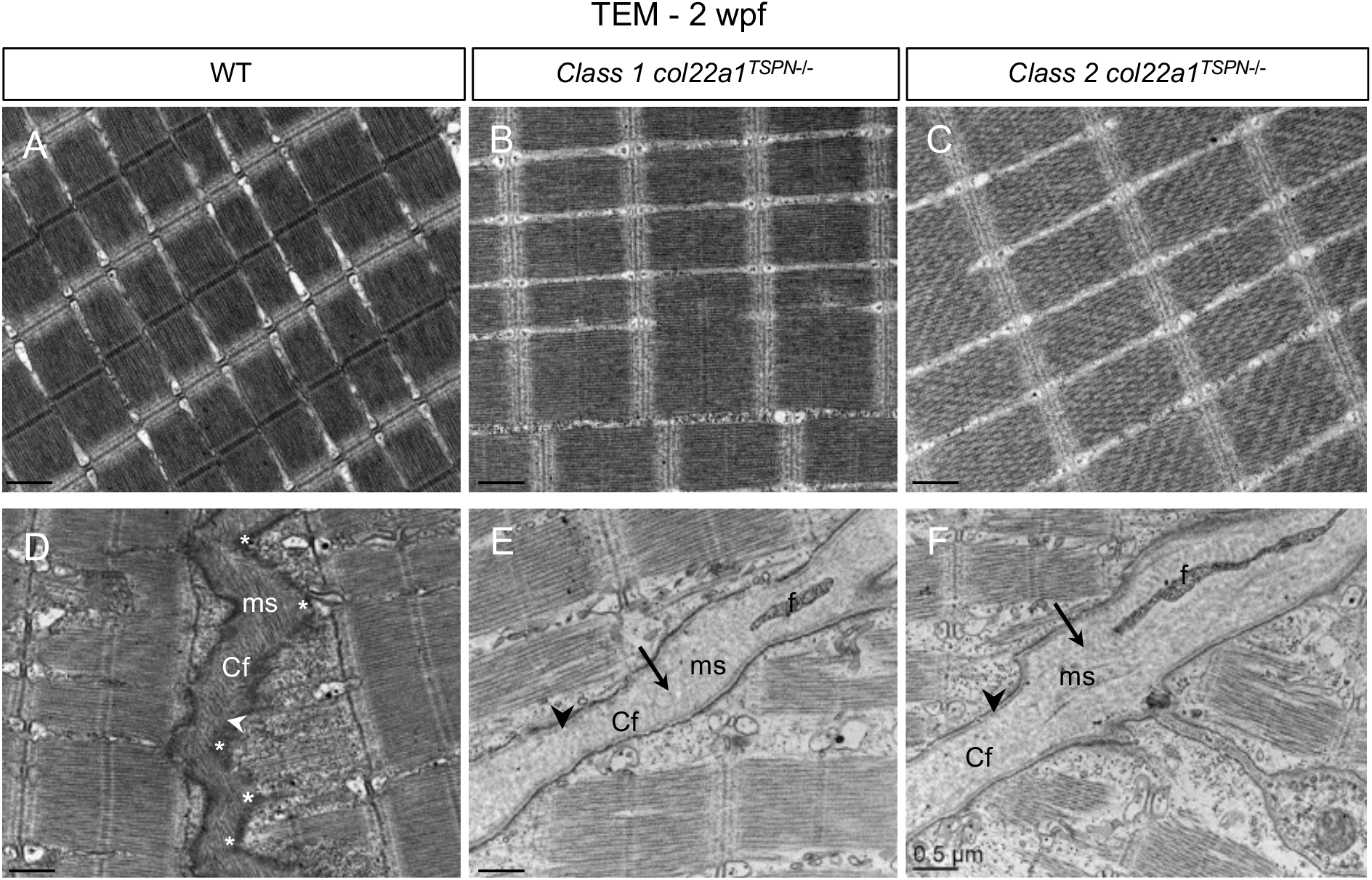
Ultrastructure of the musculotendinous unit of 2 wpf mutant larvae. Transmission electron microscopy representative images of the trunk skeletal muscle (A-C) and of the myotendinous region (D-F) of 2wpf *col22a1*^*TSPN*−/−^ class 1 (B,E) and class 2 (C,F) larvae and WT larvae (A,D). Arrows point to distended myosepta that contain sparse bundles of collagen fibrils of lucent electron density (E, F) compared to the closely packed collagen fibrils of electron density in WT myosepta (D). Arrowheads point to MTJ. The characteristic zigzag morphology of the MTJ shown in WT (D, asterisks) is not observed in class 1 (E) and class 2 (F) *col22a1*^*TSPN*−/−^ larvae. Cf, Collagen fibers; f, fibroblast; ms, myosepta; All images are longitudinal sections of the trunk muscle. All bars = 0.5μm

We then analyzed the MTJ structure and morphology of *col22a1^TSPN−/−^* larvae and WT siblings at 5dpf, when it was impossible to distinguish larvae with mild and severe phenotype. For this analysis, *col22a1^TSPN−/−^* larvae were randomly selected before being processed. Also, before morphological analysis, larvae were subjected (stimulated) or not (unstimulated) to repeated Charvet et al, (2013). Bright field observations showed that abnormally wavy muscle fibers appeared in the *col22a1^TSPN−/−^* somites after electrostimulation but not in WT (Figure 4 - Figure supplement 3A). However, somite alteration did not change birefringence measurements of the trunk regardless of whether fish are electrostimulated or not (Figure 4 - Figure supplement 3B). As observed at later stage in both phenotypical classes, an abnormal number of fibroblasts were present in *col22a1^TSPN−/−^* myosepta (Figure 4 - Figure supplement 3C) and striking reduction of sarcolemmal folds at the MTJ and distended myosepta containing sparse collagen fibrils were observed with TEM while muscle fiber ultrastructure appeared normal (Figure 4 - Figure supplement 3D). No muscle detachment was observed even after repeated contractile activity in *col22a1^TSPN−/−^* mutants (Figure 4 - Figure supplement 3D) suggestive of subtle phenotypical differences between MO-knockdown embryos and CRISPR/Cas9 mediated KO lines.

We conclude that all mutants presumably showed MTJ dystrophic phenotype with variable expressivity. In a subset of the progeny, these defects are more severe and have profound consequences on muscle fatigue due to inefficient muscle force transmission. Reduction in force transmission was obviously progressively detrimental for swimming capacity and posture, specifically in class 2 larvae.

### Lack of *col22a1* results in MTJ gene dysregulation specific to class 1 mutants

To next address the question of possible gene compensation that can explain why knock-out of *col22a1* results in variable expressivity of phenotype, we performed RT-qPCR on RNA trunk extracts of 2 wpf class 1 and class 2 mutants and WT to analyze differential gene expression of selected genes. We chose genes of MTJ components, e.g. those of the linkage systems that are likely indirect or direct binding partners of ColXXII (Charvet et al, 2013). We have previously shown a synergistic gene interaction between *col22a1* and *itga7* (Charvet et al, 2013; Figure 5A). Variation in myoseptal ECM gene expression was also sought as damaged myoseptal ECM is a common trait of class 1 and class 2 larvae (Figure 4). First, the overall results showed that expression levels of the selected genes did not follow the same pattern in class 1 and class 2 larvae (Figure 5B). Second, knockout of *col22a1* led to expression activation of the myoseptal *col1a1a* as well as *fn1b* that is only expressed at the nascent MTJ solely in class 1 larvae. This is indicative of possible remodeling of the damaged MTJ and myosepta in class 1 larvae that might not occur in class 2 larvae as these genes were even downregulated compared to WT (Figure 5B).

**Figure 5:**
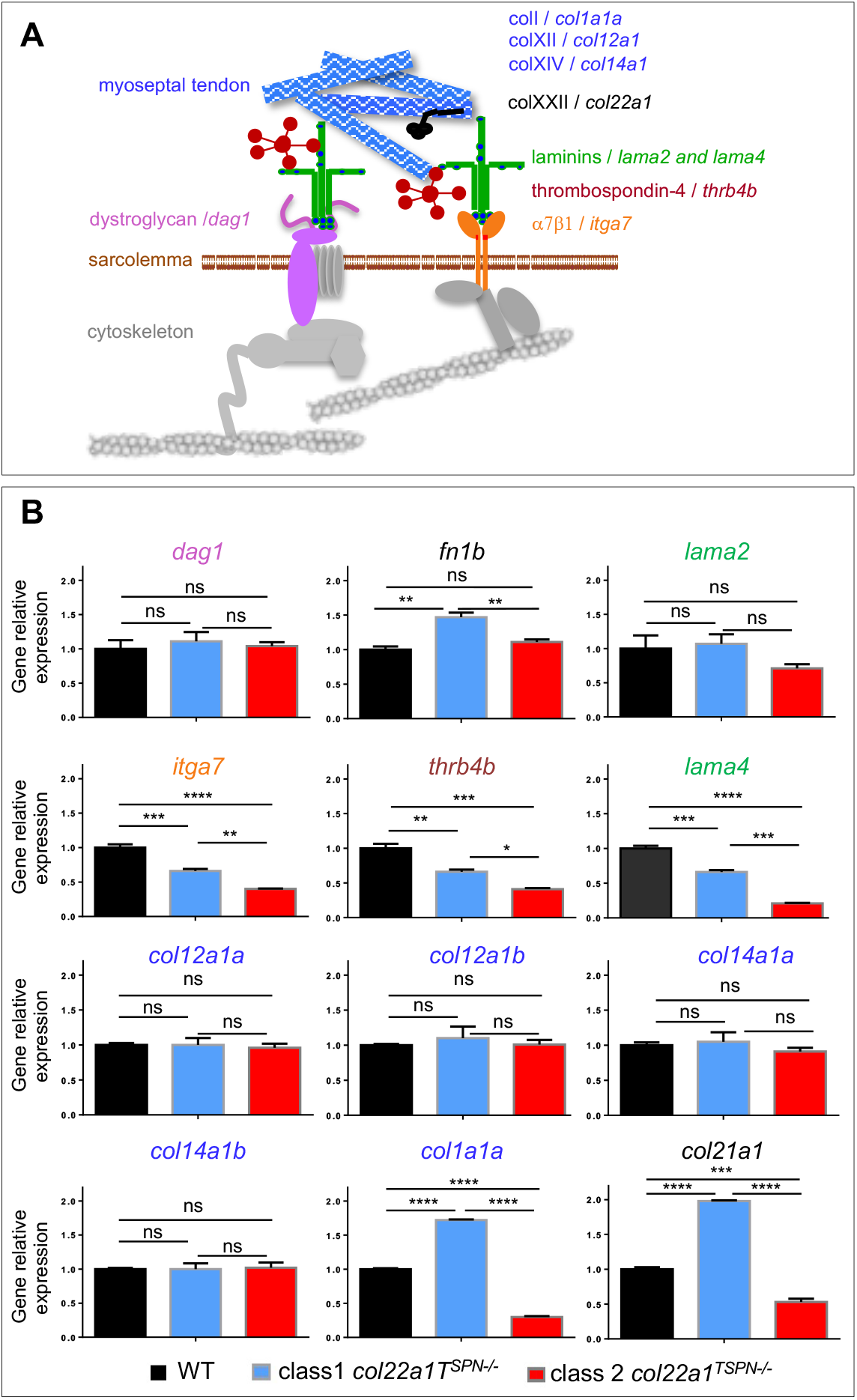
Dysregulation of the musculotendinous unit gene expression in *col22a1^−/^* mutants. A. Schematic representation of the 2 transmembrane linkage systems enriched at the MTJ, the dystroglycan-glycoprotein complex (DGC) (left) and α7β1 integrin complex (right). The presumptive location of ColXXII as part of the α7β1 integrin complex is illustrated. B. Real time RT-PCR analysis of expression of selected genes in 2 wpf trunk RNA extracts from *col22a1^TSPN−/−^* class 1 (blue) and class 2 (red) sibblings and WT (black). Data were normalized to *polr2d*. The expression levels of the different genes were compared to WT expression, which was arbitrary set to 1. Data are mean ±SEM (n=3); ****p<0.0001; ***p<0,001; **p<0,01; *p<0,05; ns, not significant.

Expression of *lama2* and *dag1* both encoding proteins of the dystrophin-glycoprotein complex (DGC) was unchanged in larvae from both phenotypic classes, as well as the myoseptal *col12a1a* and its paralog *col12a1b*, and *col14a1* paralogs for whose protein products are structurally and functionally closely related to ColXII (Bader et al, 2009, Bader et al, 2013, Nauroy et al, 2018; Nauroy et al, 2019). In contrast, expression of *itga7, lama4* and *thrb4b* was downregulated in both mutant classes compared to WT (Figure 5B). Finally, *col21a1* that is closely related to *col22a1* (Koch et al, 2004) is one of the most upregulated gene among the analyzed genes in class 1 larvae but not in class 2.

Our data suggest a common yet undescribed thread that connects the transcriptional regulation of these genes in *col22a1* mutants with mild (class 1) phenotype but not in severe (class 2) phenotype.

### Class 1 mutants MTJ phenotype is characterized by muscle fatigue and mitochondrial adaptation to exercise

We next turn to the question of why class 1 mutants do not display musculoskeletal system dysfunction. Because these mutants can reach adulthood, we have been able to measure muscle performance using a swimming tunnel respirometer that is intended to cause fish to fatigue. Fish are placed in a tunnel where water flow is increased stepwise and the capacity of the fish to swim against the current is scored (Figure 6A). U_crit_ represents the critical swimming speed originally defined as the maximal velocity a fish can reach during a swimming step protocol (Hammer 1995). In the meantime, measurements of the fish oxygen consumption during swimming tests allowed to determine the associated costs in oxygen. Two identical 170mL swimming respirometers were used in parallel and the protocol used was as depicted in Figure 6A. The tested 6mpf WT females and males showed similar U_crit_ values, therefore the mutant fish were selected for U_crit_ measurements regardless of their gender. As shown in Figure 6B, WT have no difficulty to swim against the flow at 5 Body Length.s^−1^ [BL.s^−1^]) (Figure 6B1; Video 4), and even at higher speed (6.5 to 8 BL.s^−1^) (Figure 6B2-3; Video 6). On the contrary, *col22a1^TSPN−/−^* fish swam in a steady way at a moderate speed (5BL.s^−1^) (Figure 6B4 and 7; Video 5) but were unable to sustain the effort to counter-current swimming at higher speed (6.5 to 8 BL.s-1) and their caudal fin touched several times the honeycomb in the swimming chamber (Figure 6B5-6 and 8-9; Video 7). Measurements of U_crit_ confirmed our observations as U_crit_ mean of class 1 mutant fish was statistically lower (6.98 ± 2.08 BL.s^−1^) than in WT (10.89 ± 1.05 BL.s^−1^) (Figure 6C, left panel). Similar results were obtained with *col22a1^vWA−/−^* fish (Figure 6C, right panel).

**Figure 6:**
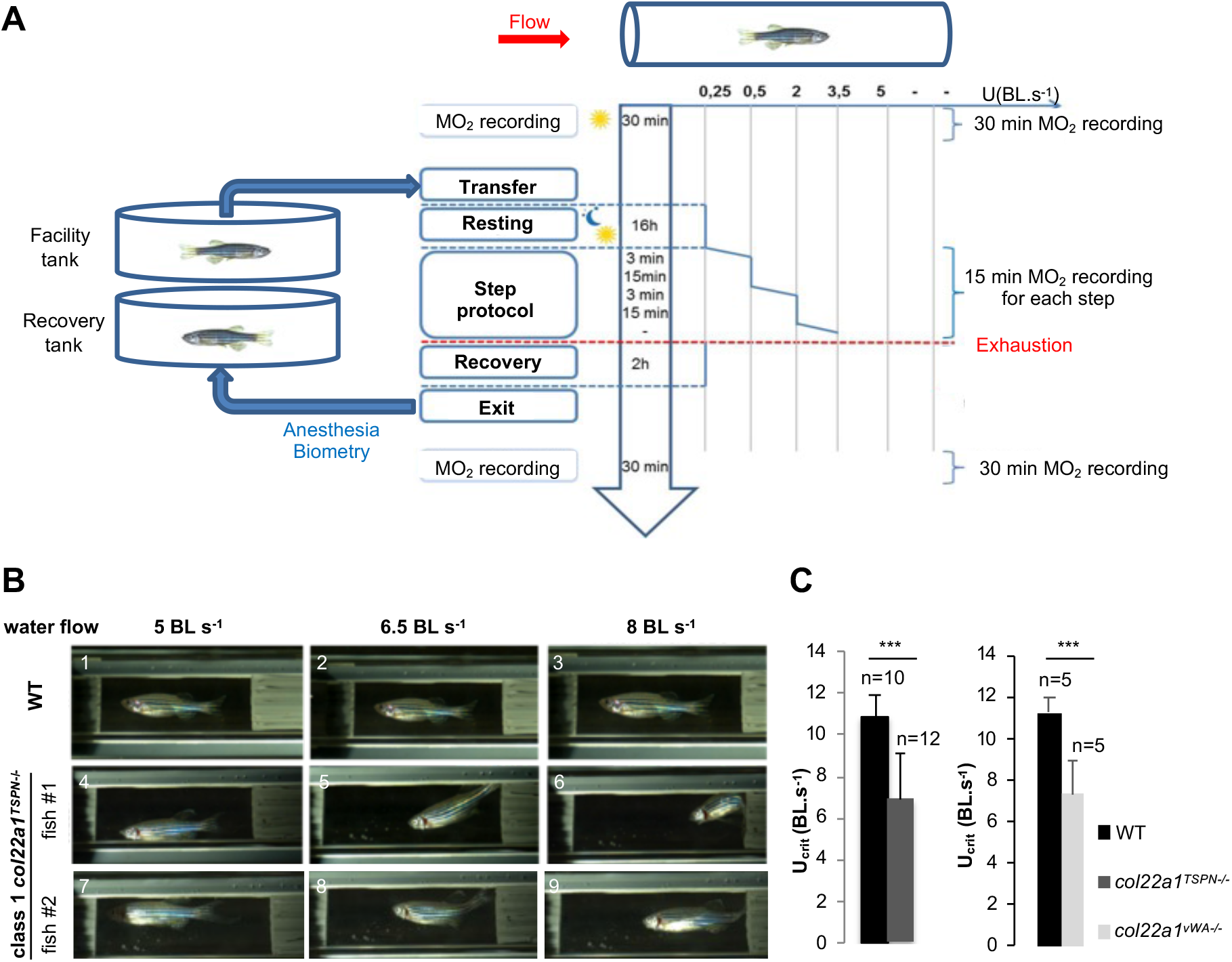
Muscle performance assays of 6 mpf *col22a1^TSPN−/−^* class 1 fish. A Flowchart of the experimental swimming step protocol tested in the swimming tunnel respirometer. B. Representative capture images of challenged *col22a1^TSPN−/−^* mutants (fish 1 and 2) and wildtype (WT) fish at different swimming speed (5BLs^−1^, 6.5BLs^−1^ and 8BLs^−1^). C Mean of critical swimming speed (U_crit_) values of (left) *col22a1^TSPN−/−^* mutants (grey bar) and WT (black bar); (right) *col22a1^vWA−/−^* mutants (grey bar) and WT (black bar). BLs^−1^, body length per second. Data are mean±SEM; ***p_value<0.001.

**Figure 7:**
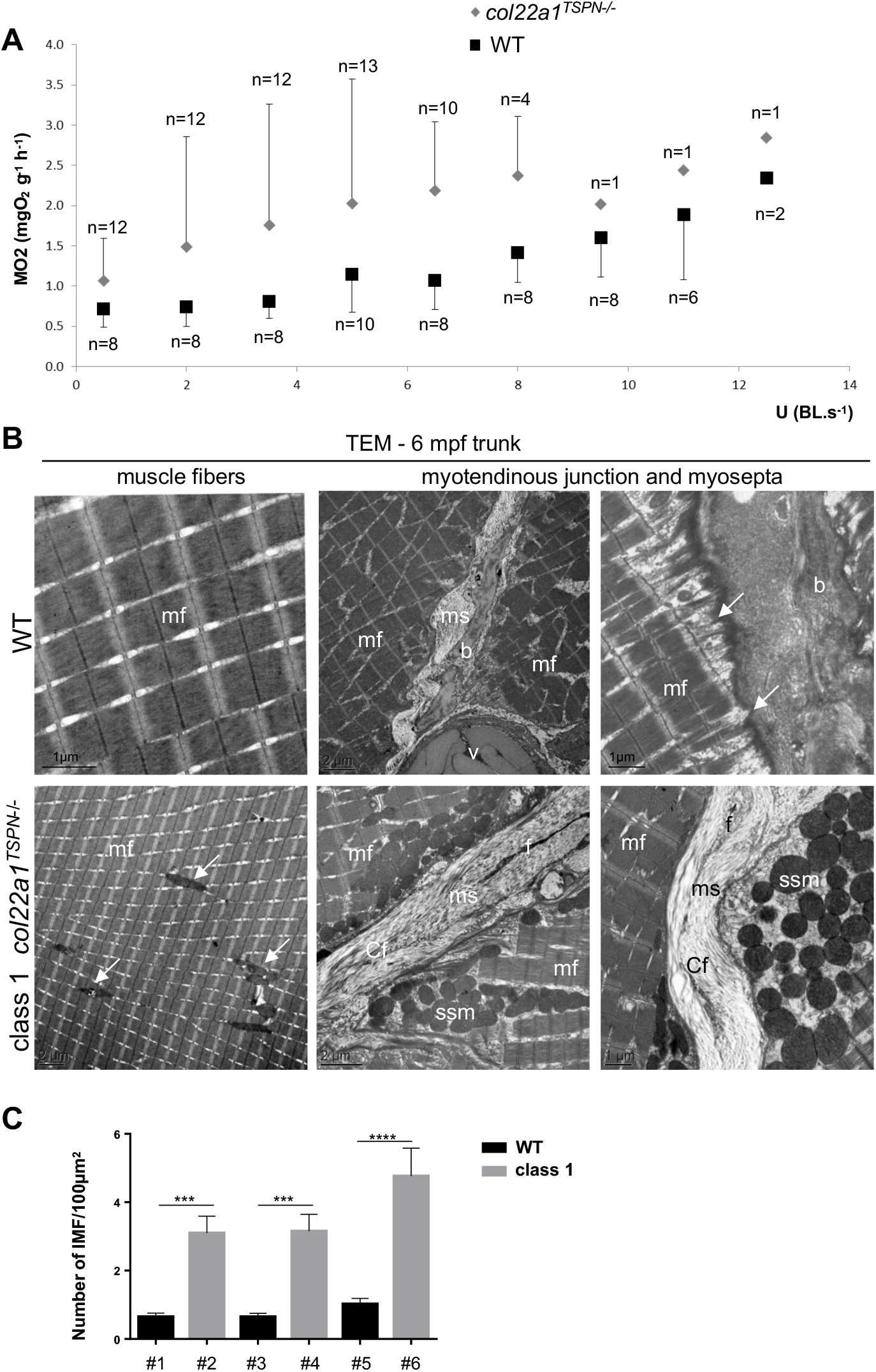
Oxygen consumption and ultrastructure of musculotendinous system of 6 mpf *col22a1^TSPN−/−^* class 1 fish. A. Measurements of oxygen consumption (MO2) during the step protocol of *col22a1^TSPN−/−^* (grey diamond) and WT (black square) fish. n, number of fish analyzed. Data are mean±SEM. B. Representative TEM images of the trunk muscle of WT (upper panel) and class 1 *col22a1^TSPN−/−^* (lower panel) fish. Upper panel: arrows point to MTJ interdigitations. Lower panel: arrows indicate IMF mitochondria. f, fibroblast; ms, myosepta; mf, myofibrils, b; bone; Cf, Collagen fibers; m, mitochondria; v, vertebra. C. IMF distribution in three different WT (#1,3,5) and *col22a1^TSPN−/−^* (#2,4,6) unchallenged fish. The total area analyzed in the three *col22a1^TSPN−/−^* and WT fish is >11,000μm^2^.

During exercise, requirements for oxygen in skeletal muscle is increased. Oxygen consumption was measured during swimming tunnel experiments and showed a clear tendency for mutant fish to consume more oxygen than WT at identical flowspeed, with a higher variability observed for mutants, though not statistically significant (Figure 7A). This could be explained by an increased trunk muscle activity in mutant fish to compensate for muscle weakness (Figure 3). Skeletal muscle mitochondria are essential to provide the energy required for movement. Resistance exercise drives skeletal muscle mitochondrial adaptations (Groennebaek and Vissing, 2017). We thus postulated that, because of the class 1 *col22a1−/−* fish muscle weakness, the continuous effort required to freely swim in the tank or even more to swim against the current results in an increased oxygen consumption that might impact mitochondrial biogenesis.

To address this question, we next examined with TEM the ultrastructure of mutant and WT trunk muscle subjected to the swimming tunnel assay (challenged fish; Figure 7B). As observed in 2 wpf mutant larvae, the most striking ultrastructural trait was the complete lack of MTJ finger-like structures in mutants compared to WT while the skeletal muscle fibers appeared normal (Figure 7B). The observation of mitochondria revealed additional differences. There are two distinct mitochondria subpopulations in skeletal muscle: the subsarcolemmal mitochondria (SS) and the intermyofibrillar mitochondria (IMF) that are found between myofibrils. They can be distinguished on the basis of their morphological features, IMF mitochondria being more elongated and thinner than SS ones (Lundbry and Jacobs, 2016). This different morphology was easily identified with TEM (Figure 7B). Interestingly, IMF mitochondria were more numerous in mutant fish compared to WT (Figure 7B). This was confirmed by quantification of IMF mitochondria in mutant and WT trunk muscle (Figure 7C). We conclude that class 1 zebrafish adult mutants have to compensate their muscle weakness due to defective contractile force transmission by increasing muscle contractile activity and metabolism probably through mitochondrial adaptation.

## Discussion

Although the MTJ is a common location for strain injuries in sports (Jakobsen and Krogsgaard, 2021), little work has focused on the interface between muscle and tendon. Therefore, how defects in MTJ integrity impacts muscle contractile force transmission have been largely overlooked. We previously shown that ColXXII MO-knockdown in developing zebrafish embryos results in the appearance of a dystrophic-like phenotype in morphants (Charvet et al, 2013). The recent development of gene editing in zebrafish has brought the use of morpholinos into question (Lawson, 2016; Kok et al, 2015). Discordance between the MO-knockdown and knockout phenotypes has been attributed to different reasons including off-target effects of morpholinos. The traits we observed in the two *col22a1* mutants we have generated here are reminiscent to those observed in 5 dpf *col22a1* morphants (Charvet et al, 2013). Nevertheless, our data unveil additional reasons for phenotypic discrepancies as possible inappropriate phenotypic analyses that prevent subtle phenotype from being revealed in genetic mutants. Also, experimenters might have not noticed the early death of developing embryos and larvae due to the large number of offspring and therefore have only analyzed the fish that display non overt phenotype.

### Lack of col22a1 results in two phenotypic classes

We found that lack of *col22a1* results in two apparent extreme phenotypic classes found in both KO zebrafish lines: *col22a1^−/−^* larva that fail to propel in swim bouts and eventually cause the *col22a1^−/−^* larva to die by starvation at around 2wpf (class 2), and *col22a1^−/−^* larva that survive to adulthood without apparent abnormalities (class 1). Nevertheless, ultrastructural analysis of the *col22a1^−/−^* mutants reveals that the two distinct phenotypical classes share common musculotendinous structural defects at the junctional extracellular matrix, though with variable grade of severity: (i) the distention of myosepta that atypically contain loosely-arranged and sparse collagen fibrils and (ii) a marked reduction of MTJ sarcolemmal folds that function is to optimize force transmission at the MTJ by increasing resistance to tensile strength. Rather, morphology of the trunk muscle and the ultrastructure of muscle fiber contractile apparatus appeared normal in both phenotypic classes. Combined ultrastructural and physiological studies revealed that ColXXII deficiency is functionally associated with an unusual MTJ phenotype. Hence, the ColXXII deficiency ultrastructural phenotype is rarely observed among others MTJ mutants or morphants. To our best knowledge, a substantial reduction of MTJ sarcolemmal folds was only reported in *itga7* knock-out mice displaying progressive muscular dystrophy (Miosge et al., 1999; Welser et al., 2009) while abnormal vertical myosepta ultrastructure with abnormal blisters within the ECM was described in the zebrafish dystrophic mutant *softy* caused by a missense mutation in *lamb2* gene (Jacoby et al, 2009). Pertaining to this observation, synergistic gene analysis in ColXXII morphants has suggested that ColXXII is part of the integrin α7β1 linkage system (Charvet et al, 2013). Here we add evidence to this assumption by showing that in absence of *col22a1, igta7* expression but not *dag1* or *lama2* is down-regulated in class 1 and 2 larvae.

### ColXXII deficiency provokes distinct defective locomotor-related behaviors

We showed that the degree of severity of the ultrastructural defects of the mutants MTJ correlates with their contractile transmission capacity as evidenced by electrically-evoked muscle contractile performance measurements and results in distinct locomotor-related behavior. The class 2 mutants displayed striking postural defects, a yet undescribed phenotype in other MTJ mutants. The capacity to correct the vertical posture is acquired at around 15-20 dpf when they begin to swim (Glasauer and Straka, 2017). At that stage, larvae are able to readjust their posture by swimming bouts, a way to fight against gravity, and to this end, larvae control the speed and length of muscle contractions. Zebrafish, as all teleosts, have a denser head than tail and thereby they have to correct the destabilizing nose-down rotations to swim (Ehrlich and Schoppik, 2017). Maintenance of posture requires continuous active sensorimotor control. Our findings revealed timely unexpected functional consequences of the MTJ ultrastructural defects. In *col22a1^−/−^* larvae, postural behavior is defective secondary to ineffective muscle force transmission. As such, muscle contractile activity is reduced, swimming bouts are ineffective and the vertical posture cannot be fought. This in turn injured the chances of *col22a1^−/−^* larvae to swim and search for food resulting in starving to death, this latter assumption being supported by the empty-gut phenotype of class 2 larvae.

Contrary to class 2 mutants, the structural defects in the musculotendinous unit did not seem to affect dramatically class 1 larval motility, swimming capacity or posture although electrically-evoked contractile force measurements of the trunk musculature revealed muscle weakness. Class 1 adult fish had to be subjected to swimming-induced exercise to reveal the functional consequences of *col22a1* gene ablation in zebrafish. We showed that these fish rapidly developed muscle fatigue and reduced swimming capacity due to defective force transmission capacity resulting from MTJ ultrastructural defects. O_2_ consumption was substantially increased in class 1 mutants during swimming performance assays. One plausible explanation is that during sustained swimming activity, *col22a1^−/−^* zebrafish have to initiate skeletal muscle contractions more often for a same effort to compensate for muscle weakness. This likely entailed a higher energy demand. This is consistent with the trend of increased oxygen demand observed during the swimming challenge. Although not significant, class 1 mutant adults indeed consumed more oxygen than WT for a same effort. Along this line, a higher number of intermyofibrillar mitochondria (IMF) was observed in class 1 muscle fibers. Acute increased in contractile activity patterns, as during exercise training (endurance or intermittent), was reported to favor mitochondria biogenesis (Hood et al., 2016). The daily swimming activity of class 1 fish might require an increased energy demand as the force transmission capacity is defective in these mutants, that can provoke a mitochondrial adaptation over the long term.

In addition, IMF have been associated to energy production for contractile activity (Ferreira et al, 2010). In that, our data suggested a mastryoshka doll-like mechanism by which defective MTJ activity resulted in useless force transmission that could subsequently increase muscle contraction frequency, which in turn augment the overall muscle energy demand that led to training-induced mitochondrial adaptation.

### Why lack of col22a1 gene produces phenotype with variable severity?

Our finding raised the question of why lack of *col22a1* gene produces different phenotypic patterns. The relationship between genotype and phenotype is not simple particularly in ECM genes (Pereira et al, 1994; Jin et al, 2007; Li et al 2008). This can be due to incomplete penetrance or variable expressivity, two events often observed in human inherited connective tissue disorders (Lobo, 2008). This was also reported as a possible mechanism to explain the reported phenotypic discrepancies in zebrafish, specifically when mutant phenotype was compared to the morphant one (El-Brolosy and Stainier, 2017; Kontarakis and Stainier, 2020). As discussed above, in our study the phenotype of our CRISPR/Cas9 *col22a1* mutant lines mostly recapitulates the one reported for *col22a1* morphants that also display gradient of phenotypic severity (Charvet et al, 2013), these phenomena may account for the surprising apparent lack of phenotype in class 2 mutants. But gene compensation can also be invoked to explain this discrepancy (Rossi et al., 2015). In support to this assumption, differential dysregulation of MTJ gene expression occurred in class 1 but not class 2 mutants. A significant activation of *col21a1*, a *col22a1* closely-related gene and *col1a1a* are observed in class 2 but not class 1 larvae. *col21a1* gene has the particularity to be present in zebrafish and human genomes, but not in rodent genomes (Fitzgerald and Bateman, 2004). The function of this gene is still unknown. *col21a1* transcript is expressed in developing fish but its expression pattern is still unclear (Malbouyres M and Ruggiero F, personal communication, 03/2020). Instead, the upregulation of *col1a1a* in class 2 mutants can be interpreted as a way to compensate the reduced density of collagen fibrils in mutant myosepta and maintain to some extent the MJT activity. In support to this assumption, a high number of fibroblasts, the main cell source of collagen I, was abnormally observed in *col22a1* mutants, though independently on the phenotypic classes. Nevertheless, several points remain obscure. Specifically, we cannot explain the severity of the *col22a1* phenotype in 30% of the larvae while about 70% display a mild phenotype. This will require further investigation including the search for expression of gene modifiers as recently reviewed in (Rahit and Tarailo-Graovac, 2020).

In conclusion, the musculotendinous disorder described here is unusual in that it is not associated with defects in muscle fiber ultrastructure and/or alteration of the contractile unit function but with useless force transmission that results in posture and locomotion disabilities.

In that respect, *COL22A1* should be viewed as a candidate gene for human muscular dystrophies with unresolved genetic cause. Our study also provides a valuable genetic tool to investigate variable expressivity often observed in connective tissue human diseases and that represents an important point to consider for the design of chemical screens.

## Materials and Methods

### Zebrafish strain, maintenance, specific treatments and ethics statement

Zebrafish (AB/TU) were raised and bred according to standard procedures (Bader et al., 2009) (PRECI, SFR Biosciences UAR3444/CNRS, US8/Inserm, ENS de Lyon, UCBL; agreement number C693870602). The developmental stages are given in hours (hpf), days (dpf), weeks (wpf) and months (mpf) post-fertilization at 28°C, according to morphological criteria. For optimal growth during the first ten days, the embryos (30 individuals per 300 ml) are kept in tanks filled with water at least 5 cm deep and raised at 28.5°C. Then, the water volume is progressively increased to 900mL and the water circuit is open. At 6dpf, embryos are fed three times a day (Gemma75). From 12dpf, embryos are fed with live brine shrimp every day. Tricaine (ms-222, Sigma-Aldrich, St Louis, Missouri, USA) was used to anesthetize fish. From 24hpf, embryos were treated with phenylthiourea (P7629, Sigma-Aldrich, St Louis, Missouri, USA) to prevent pigmentation. All animal manipulations were performed in agreement with EU Directive 2010/63/EU. The creation, maintenance and use of *col22a1*^TSPN−/−^ and *col22a1*^vWA-^ lines were approved by the local ethics committee (CECCAPP_IGFL_2014_002 and CECCAPP_PRECI_2017_003) and all procedures have been approved by the French ethical committee (APAFIS#13345-2017113012206024v3).

### Generation of CRISPR/Cas9 KO lines

DR274 gRNA plasmids and Cas 9 protein were from Addgene (Cambridge Massachussets, USA) and New England Biolabs (Ipswich, MA, USA), respectively. CRISPR gRNAs were designed using CRISPR design software (http://crispor.tefor.net/ and http://crispr.dbcls.jp/). Different sgRNAs have been tested and we selected the two with the highest mutagenesis efficiencies. The DNA binding sites targeted by these sgRNAs are localized in the vWA domain (5’-GGATAAGACACGTGTGGCAG-3’) and in the TSPN domain (5’-GGATGGCGAGAACAGGGCGG-3’). Oligonucleotides were annealed in a thermo block at 95°C for 5 min followed by a slow cooling to room temperature (≈25°C) and cloned in DR274 gRNA plasmid between BsaI sites. All constructs were verified by sequencing. To make gRNA, the template DNA was linearized by DraI (NEB-R0129) digestion and purified using a NucleoSpin Gel and PCR Clean-up (#740609, Macherey-Nagel, Duren, Germany) kit. gRNA was generated by *in vitro* transcription using a T7 MEGAscript transcription kit (AM1334-Thermo Fisher Scientific, Waltham, Massachussets, USA). After *in vitro* transcription, the gRNA was purified using ammonium acetate precipitation and stored at −80°C. Cas9 protein (2.12 ng) (New England BioLabs Inc. Ipswich, MA, USA,) and gRNA (133 pg) were co-injected in the cell at the one-cell stage and a quarter of the lay was used to evaluate mutagenesis efficiency by HRMA as previously described in (Talbot and Amacher, 2014).

### Genotyping

For gDNA extraction, embryos or fin clip were placed in PCR tubes in 200μL of NaOH 50mM, incubated 10 minutes at 95°C and immediately placed on ice. 20μL of Tris-HCL 1M pH8 were added to buffer the solution and gDNA were stored at 4°C. The primers used to genotype *col22a1^vWA^ and col22a1^TSPN^* mutants were designed using Primer 3 Plus software (www.bioinformatics.nl/cgi-bin/primer3plus/primer3plus.cgi), genotyping : F-5’CCAGGTTGTAAGAACGTCCAC3’/R-5’GAACAACCTGATCCCAGCAG3’ (for *col22a1^vWA^*) and F-5’CATCCCGTAAGGAAGACTGG3’/ R-5’GATGGGTAGCGTCTCGATGT3’ (for *col22a1^TSPN^*). The amplicons for each line were run simultaneously in a 2% agarose gel and were sequenced.

### RNA extraction and quantitative real-time PCR

Total RNA was isolated using Trizol (Invitrogen, Waltham, Massachussets, USA). Reverse transcription was performed using M-MLV reverse transcriptase (Promega, Fitchburg, Wisconsin, USA) for 1h at 37°C in presence of random hexamers (Promega). qPCR primers (supplement Table 1) were designed to produce amplicons of approximately 200 bp. qPCR was performed using a SYBR Green mix and fast amplicon protocol according to manufacturer instructions (Roche, Penzberg, Germany). Data were analyzed using the Ct method and normalized to Pol II. Student’s t-tests with Welch’s corrections were used to determine statistical significance (n=3).

### Immunofluorescence staining

Whole-mount staining was performed as described previously (Bader et al., 2009). Wholemount embryos were observed with a Zeiss LSM 780 spectral confocal microscope (Zeiss, Oberkochen, Germany). The primary and secondary antibodies used in this study are listed in (supplement Table 2). Nuclei were stained with Hoechst solution and F-actin with fluorescent phalloidin conjugates (all from Sigma-Aldrich, St Louis, Missouri, USA)

### Birefringence assay

WT or *col22a1*^TSPN−/−^ 5 dpf larvae were anesthetized with tricain, placed on a glass slide between two polarizers (3Dlens, Taiwan) and observed with a Zeiss Axiozoom V16 equipped with a AxioCam MRm 1.4 megapixels camera before and after electrostimulation. The mean gray value of pixels in the trunk region was measured with ImageJ software. Values are expressed as the percentage of unstimulated WT larvae +/- SD (n=10). Data were analyzed by one-way analysis of variance (ANOVA) followed by Brown-Forsythe and Bartlett’s multiple comparison tests (GraphPad Prism software).

### Transmission electron microscopy (TEM)

Zebrafish embryos and larvae were fixed in 1.5% (v/v) glutaraldehyde and 1.5% (v/v) paraformaldehyde in 0.1 M cacodylate buffer (pH 7.4) several days at 4°C followed by washing in 0.1 M cacodylate buffer, 8% sucrose (10 minutes, 3 times). Then, samples were quickly washed in distilled water. After post-fixation in 1% osmium tetroxide in 0.1 M cacodylate buffer, 8% sucrose for 45 minutes at room temperature, samples were dehydrated in graded series of ethanol (30 to 100 %), 5 minutes each, and propylene oxide during 10 minutes. Finally, samples were embedded in epoxy resin. Ultrathin sections were stained with 7% uranyl acetate in methanol and lead citrate, and observed with a Philips CM120 electron microscope (Centre Technologique des Microstructures, Université Lyon 1, France) equipped with a Gatan Orius 200 2Kx2K camera.

For adult muscle tissues, 6-months trunk muscles were dissected in the fixative buffer under stereoscopic control and skin was delicately removed thanks to tweezers. Those thicker samples were treated as described above but with an extended sample incubation time (15 mins for each step). An additional 15 mins incubation in propylen oxide/ethanol 100% (v/v) followed by 2 X15 mins incubation in propylen oxide were performed. Times of incubation during the embedding steps were also prolonged.

### Muscle force measurements

Procedures and measurements were performed as previously reported in (Charvet et al, 2013). The detailed procedure and changes to the previous protocol are described in Supplementary Material and Methods.

### Swimming performance assays and oxygen consumption measurements

Swimming performance tests and oxygen consumption measurements were performed as previously described in (Lucas et al., 2016). The detailed procedure and measurements are detailed in Supplementary Material and Methods. Videos were acquired during swimming step protocol experiments, at a moderate (5BL.s^−1^) and quite rapid (8 BL.s^−1^) water flow respectively. Movies are played 10 times slower than original speed. Biometry of the fish used for these experiments is shown in (supplement Table 3). Swimming performance assays was performed in both lines and oxygen consumption was only measured in *col22a1^TSPN−/−^*. Live 2 wpf larvae were placed in Danieau media. The motility of larvae was recorded using a Zeiss Axiozoom V16 stereoscopic microscope with a high resolution AxioCam HRm Rev.3 Firewire camera. To assess swimming activity of the fish, video-recorded at 5 ms/frame, during 25s was performed without any stimulation. Videos of 6mpf zebrafish were recorded using a high-speed camera MiroM310.

### Statistical analysis

Statistical analyses were carried out using GraphPad Prism software. Depending on experiments, Student’s t-tests or Anova were used to compare WT and *col22a1^−/−^* fish as indicated. Tukey multiple comparison tests were used for oxygen consumption measurement statistical analyses,

## Acknowledgments

We thank Prof Manuel Koch (University of Cologne) and Dr Loïc Teulier (LENHA, Lyon) for helpful discussion. We acknowledge the contribution of the zebrafish facility (PRECI, SFR Biosciences-Gerland), specifically Robert Renard for his assistance with the swimming tunnel implementation, Cherif Kabir for his helpful IT assistance with the swimming tunnel software, Loup Plantevin (INSA, Lyon) for his kind help with the implementation of the swimming tunnel calibration. We thank the ‘Centre Technique des Microstructures’ (UCBL, Villeurbanne) for technical assistance with transmission electron microscopy. This work was supported by the CNRS and the “Association Française contre les Myopathies” [MNM1-2010] to FR. AG is a recipient of a post-doc fellowship from the “Association Française contre les Myopathies”. PN is a recipient of the French government (NMRT) and the “Fondation pour la Recherche Médicale” (FDT20160435169) fellowships.

## Author contribution

Conceptualization: MM, AG, BA, CL, FR - Investigation: MM, AG, BA, FR - Methodology: MM, AG, BA, CL, LB, MS, PN, FS - Visualization: MM, FR - Funding acquisition: FR - Writing: Original draft: FR, MM - Review and editing: MM, AG, CL, FS, BA, FR.

