## Supplemental File for "The myotendinous junction marker collagen XXII enables zebrafish postural control learning and optimal swimming performance through its force transmission activity"

### Supplementary Materials and Methods

*Muscle contraction measurements*

Procedures and measurements were performed as previously reported (Charvet et al, 2013). Larvae are briefly placed in a beaker containing 0.168 mg/mL of tricain (3-amino benzoic acidethylester, Sigma-Aldrich, A5040). The head is then crushed and larvae are transferred to a Tyrode physiological solution containing (in mM) 140 mm NaCl, 5 mm KCl, 2.5 mm $CaCl_2$, 1 mm $MgCl_2$, 10 mm Hepes, pH 7.2. Under binocular control, fish head and tail extremities were glued with surgical glue (Histoacryl, B Braun) on homemade thin aluminium foils pierced with a small hole allowing to attach the head portion of the fish to the arm of an AE801 force transducer and the tail portion to a fixed pin. Fish were stimulated by field electrodes placed on either side of the fish using a Harvard apparatus 6002 stimulator. In each fish, the voltage of 0.5 ms duration pulses was increased to give maximal twitch response and the voltage used for measuring maximal force was set at 20 % above this maximal value. The fish was then gradually stretched on the side of the transducer moved with a micromanipulator until maximal twitch force was obtained. The force signal was recorded at 10 kHz sampling frequency using the WinWCP software (Strathclyde University, UK) driving an AD converter (National Instruments, USA).

*Swimming performance tests and oxygen consumption measurements*

Swimming performance tests and oxygen consumption measurements were performed as previously described (Lucas et al., 2016). Critical swimming speed (i.e. the maximal velocity a fish can reach during a swimming step protocol, $U_{crit}$; Hammer, 1995) is frequently employed as an indicator of swimming capacities. Since $U_{crit}$ notably depends on the maximal ability of fish to provide energy during sustained swimming activity, these studies are often associated with the assessment of metabolic performance. Aerobic metabolic scope (AMS; Brett, 1964; Fry, 1971) is defined as the difference between the active metabolic rate (AMR), which is the highest metabolic rate the organism can sustain under maximal activity and the standard metabolic rate (SMR), the metabolic rate necessary to maintain vital functions and measured under resting conditions.

Experimental set-up - Two identical 170 mL swimming respirometers (Loligo Systems, Denmark) were used to assess the swimming and metabolic performance of fish. Each swimming respirometer was composed of (a) a swimming chamber, where the fish was placed to be tested, (b) a motor fitted with a three-bladed propeller to control water flow and (c) honeycomb placed at each side of the swimming chamber to laminarize the water flow. Each swimming respirometer was submerged in a 20 L buffer tank, and filled with temperature controlled (i.e. 28°C) and oxygenated mixed water as in the rearing system.

The oxygen consumption ($MO_2$ in mg $O_2$ $g^{-1}h^{-1}$) associated with the activity of fish was measured by intermittent flow respirometry. Water supply in each swimming respirometer was provided by flush pumps, which controlled water flow from the buffer tank to the swimming respirometer. This allowed alternation between phases of oxygen renewal and phases of $MO_2$ measurement with a cycle of 3: 15 min. An oxygen probe (fibre optic sensor, PreSens, Germany) was used to record oxygen concentration in the swimming respirometer where the fish was placed. The probe was connected to a multichannel oxygen measuring system (OXY 4 mini, PreSens, Germany) to record the level of dissolved oxygen in the water every 5 s. Optic fibers was calibrated once at the beginning of the swim test using 0% and 100% air saturation for a controlled temperature of 28°C. A conversion factor based on oxygen solubility into water was used to convert oxygen data from per cent saturation to $mgO_2L^{-1}$ (*i.e.* 100% was equivalent to 7.94 $mgO_2$ $L^{-1}$ for a 28°C temperature and 0 salinity).

Experimental procedure –The fish were starved the morning before being transferred individually (late afternoon) to one of the two swimming respirometers for 24hrs before the swimming challenge. Each experimental trial consisted of challenging two fish in parallel. Each individual fish was left undisturbed with a water flow of 0.25 BL $s^{-1}$ for a night to permit recovery from handling stress and acclimate to this new environment. The swimming challenge started by increasing water flow by steps of 1.5 BL $s^{-1}$ from 0.5 to 12.5 BL $s^{-1}$. Each step lasted 15 min, during which fish $MO_2$ was measured. Note that between 2 steps, $O_2$ is renewed; and that each time, it takes 3 minutes to progressively increase the water flow to reach the next step. The experiment stopped when the fish fatigued, i.e. when it did not manage to swim against the current and stay on the honeycomb. Speed was then decreased to 0.25 BL $s^{-1}$ for a recovery period of 2 h. Fish were then removed from the swimming respirometer and anaesthetized using tricaine. Zebrafish are placed into a beaker containing 0.168 mg/mL of tricaine (3-amino benzoic acidethylester, Sigma-Aldrich, A5040, stock solution at 4 mg/mL dissolved in distiller water and stored at -20°C) freshly dissolved in system water. Fish progressively stop moving and we consider that they are anesthetized when they stop reacting to an external stimulation (e.g. gently taping on the bench close to the beaker). Standard and total length, mass and sex of each individual were determined; characteristics of the fish tested in the swim tunnel are described in Table S3. Before and after each trial, a blank measurement was performed to quantify microbial oxygen consumption in the swimming respirometer. The average of these two values was subtracted from the measured oxygen consumption. After each individual test, equipment was fully cleaned to reduce microorganism development. Each fish was tested once. Note that during swimming challenge experiment, fish are maintained in the same water conditions (temperature, pH, osmolarity…) than in their original animal facility.

Critical swimming speed - The critical swimming speed $U_{crit}$ was calculated according to the formula of Brett (1964):

$U_{crit} = U_t + t_1 * t^{-1} * U_1$

where $U_t$ (in BL $s^{-1}$) is the highest velocity maintained for an entire step, $t_1$ (in min) is the time spent until the exhaustion of fish at the last step, t (in min) is the swimming period for each step (i.e. 15 min in the present study) and $U_1$ is the increment velocity (1.5 BL $s^{-1}$).

*Oxygen consumption ($MO_2$) measurements* - Oxygen consumption $MO_2$ is expressed in mg $O_2$ $g^{-1}$ $h^{-1}$ and calculated using the following formula:

$MO_{2meas} = \Delta [O_2]. V. \Delta t^{-1} . M_{meas}^{-1}$

where Δ[O2] (in mg $O_2$ $L^{-1}$) is the variation in oxygen concentration during the measurement period Δt (in h), V (in L) is the volume of the respirometer minus the volume of the fish and $M_{meas}$ (in g) is the fish mass measured. An allometric relationship exists between oxygen consumption and body mass, which permits correction of $MO_{2meas}$ using the following formula:

$MO_{2cor} = MO_{2meas.} (MO_{2meas}. M_{cor}^{-1})^{1-b}$

where $MO_{2cor}$ (in mg $O_2$ $g^{-1}$ $h^{-1}$) is the oxygen consumption related to a standard fish of 0.1 g ($M_{cor}$), $MO_{2meas}$ (in mg $O_2$ $g^{-1}$ $h^{-1}$) is the oxygen consumption estimated for experimental fish whose mass was $M_{meas}$ (in g) and b is the allometric scaling exponent describing the relationship between oxygen consumption and body mass was calculated according to Lucas et al. (2014).

### Supplementary Figures

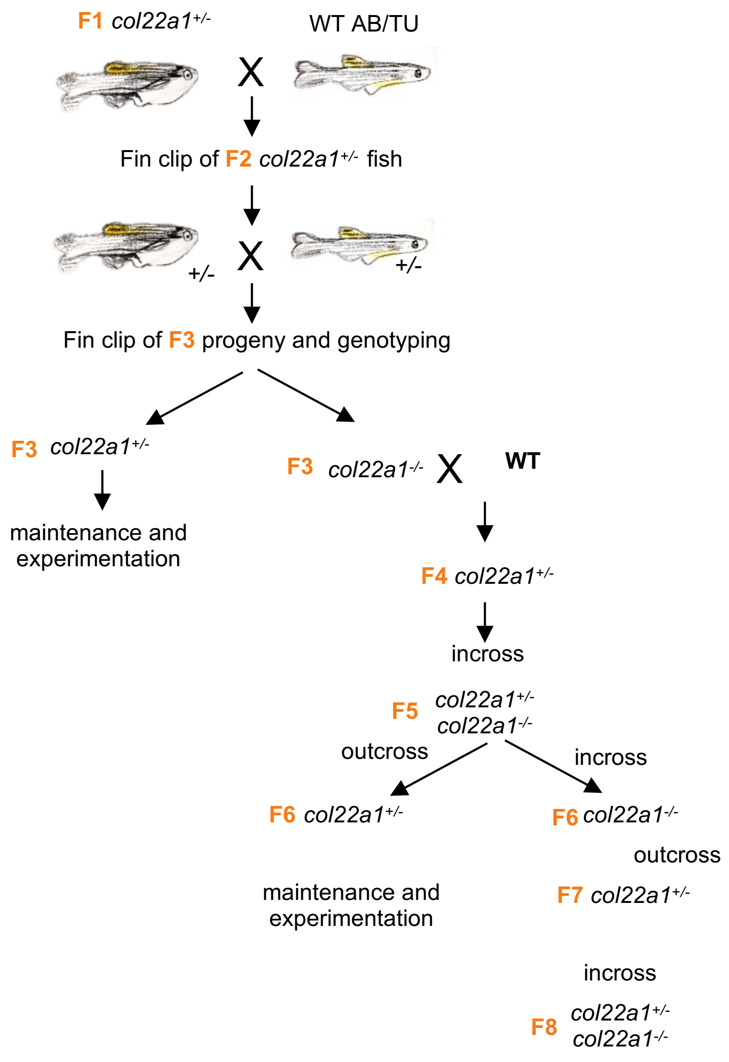


**Figure S1: CRISPR-Cas9 *col22a1* lines -** generation, crossing and maintenance. The same protocol has been applied to the two mutant lines (*col22a1*$^{vWA}$ and *col22a1*$^{TSPN}$).

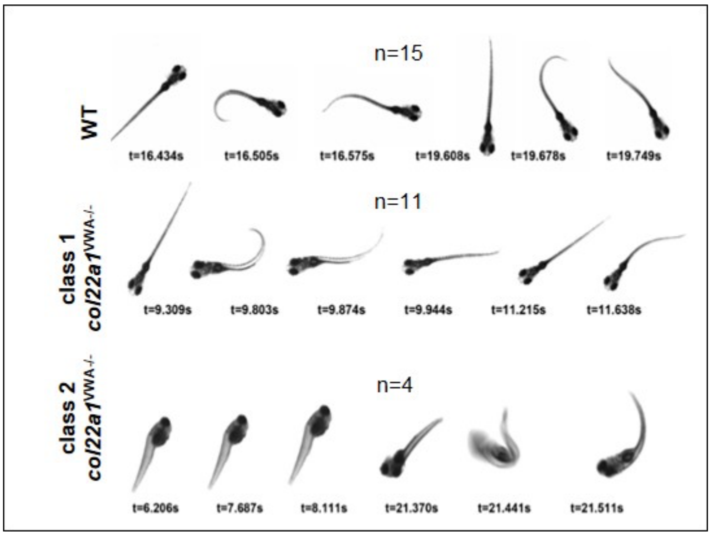


**Figure S2: Postural and swimming behaviors of 2wpf larvae.** Single frame from videos of 2 wpf *col22a1*$^{vWA-/-}$ class 1 (row 2) and class 2 (row 3) larvae and WT (row 1). t, time expressed in seconds (s). n= number of analyzed larvae. wpf, weeks post-fertilization

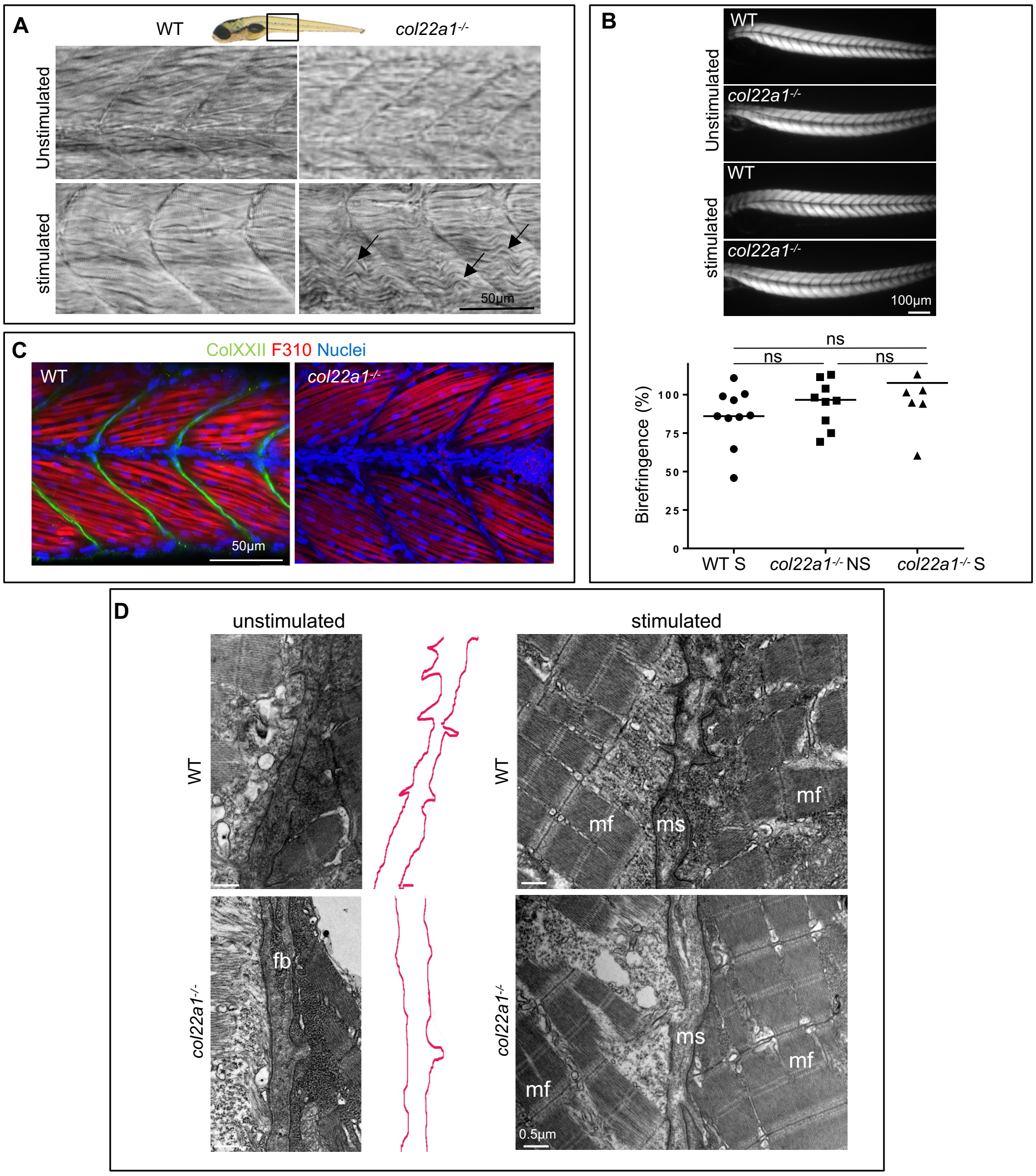


**Figure S3: Structural analysis of *col22a1*$^{TSPN-/-}$ trunks at 5 dpf.** A. Representative bright field images of *col22a1*$^{TSPN-/-}$ and wildtype (WT) larvae before (upper) and after repeated electric stimulation (lower). Arrows point to abnormally wavy muscle fibers. B. Birefringence images of *col22a1*$^{TSPN-/-}$ and WT larvae before (upper) and after (lower)r epeated electric stimulations. Quantification: values represent mean normalized to unstimulated WT. NS, unstimulated; S, stimulated. C. Whole-mount immunofluorescence staining with antibodies against ColXXII (green) and the myosin light chain 1 marker (F310, red). Nuclei are in blue. An abnormal high number of fibroblasts are observed in *col22a1*$^{TSPN-/-}$ specimen. Anterior is towards the left. D. TEM images of the trunk of *col22a1*$^{TSPN-/-}$ and WT embryos before (left panel) and after (right panel) electrostimulation. Red lines delineate the MTJ. Note the reduction of sarcolemmal folds and distended myosepta with sparse collagen fibrils in *col22a1*$^{TSPN-/-}$ specimen. fb, fibroblast; mf, muscle fiber; ms, myosepta. All bars=0.5 µm.

**Supplementary Tables**

**Table S1:** List of primers. For col22a1, the domains in which the primer sequences have been designed are indicated into brackets: COL5, collagenous domain 5; TSPN, Thrombospondin-1 N terminal-like domain; vWA, von Willebrand-like domain.

| Gene | Primers |
|---|---|
| *col1a1* | *F-5'CTGCAAGAACAGCATTGCAT3'*<br>*R-5'TAGGCAGACGGGATGTTTTC3'* |
| *col12a1a* | *F-5'AGGGCTCGTCTGTGTCTGAT3'*<br>*R-5'GTTTTGCAGGATGACCGAGT3'* |
| *col12a1b* | *F-5'CTCCTCAGGACAAGGAGCAC3'*<br>*R-5'GCACCAGCTTTTCTCCAGAC3'* |
| *col14a1a* | *F-5'TGCTCATTTCTGAGGTGACG3'*<br>*R-5'GCGAACACTGCAATCTCGTA3'* |
| *col14a1b* | *F-5'GACCACGCTTCCTCTGACTC3'*<br>*R-5'AGGTTACGCAGCTCCACTGT3'* |
| *col21a1* | *F-5'CTGGACCTGAGGGACGAC3'*<br>*R-5'CCGATCGGAGGATGTCTCT3'* |
| *col22a1(*vWA*)* | *F-5'CAAGTGTGGGCAAGGAGAAT3'*<br>*R-5'CGTTTGACCTCCTCCAATGT3'* |
| *col22a1 (*COL5*)* | *F-5'AGATGGAGCGGACGGTTT 3'*<br>*R-5'CTGGTGGGCCTTCATCTC3'* |
| *col22a1* (TSPN*)* | *F-5'CATCCAGGGGAAAACTGTCA3'*<br>*R-5'CCATCCTCTGCAGGTCAAAG3'* |
| *dag1* | *F-5'TGTTGGAGCCTCTGTTACCC3'*<br>*R-5'TGGGTTTCTTTGGAGGTCTCT3'* |
| *fn1b* | *F-5'ATTCAACGCACGTTCCTACC3'*<br>*R-5'TAATTTTGCCCTTGCCTGAC3'* |
| *itga7* | *F-5'CAGCTGGTACCCAAACACG3'*<br>*R-5'CACAGAGTACCCGAATGAGGA3'* |
| *lama2* | *F-5'CCACTGGGGTCACGACAT3'*<br>*R-5'CGGTGTTCAGGTTACACTGC3'* |
| *lama4* | *F-5'AATCACGTTGTGGGGTCTGT3'*<br>*R-5'ACCGATAAACACTGGCGAAC 3'* |
| *polr2d* | *F-5'CCAGATTCAGCCGCTTCAAG3'*<br>*R-5'CAAACTGGGAATGAGGGCTT3'* |
| *thrb4b* | *F-5'GAGGAGGAAGACTGGGAGTGT3'*<br>*R-5'GTCCTCTGGGATGGTGTCAT3'* |

**Table S2:** List of primary (in bold) and secondary antibodies used in this study.

| Primary and Secondary antibodies | Reference | Dilution used |
|---|---|---|
| **anti-ColXXII** | polyclonal, rabbit (home-made[1]) | 1:1000 |
| **anti-ColXII** | polyclonal, guinea pig (home-made[2]) | 1:250 |
| **F310, myosin light chain 1** | DSHB, F310, monoclonal, mouse | 1:20 |
| anti-rabbit Alexa Fluor 488 | Invitrogen, A11034 | 1:500 |
| anti-mouse Alexa Fluor 488 | Invitrogen, A11029 | 1:500 |
| anti-guinea pig Alexa Fluor 488 | Invitrogen, A11073 | 1:500 |
| anti-rabbit Alexa Fluor 546 | Invitrogen, A11035 | 1:500 |

[1]Charvet B, Guiraud A, Malbouyres M, Zwolanek D, Guillon E, Bretaud S, Monnot C, Schulze J, Bader HL, Allard B, Koch M, Ruggiero F. 2013 Knockdown of col22a1 gene in zebrafish induces a muscular dystrophy by disruption of the myotendinous junction. Development, 140:4602-13.
[2]Bader HL, Charvet B, Veit G, Driever W, Koch M and Ruggiero F. 2009 Zebrafish collagen XII is present in embryonic connective tissue sheaths (fascia) and basement membranes. Matrix Biol 28:32-43 *doi*: 10.1016/j.matbio.2008.09.580

**Table S3:** Biometry of fish used for swimming tunnel step protocol. M, male; F, female; SL, standard length; TL, total length.

| Fish genotype and number | Sex | Weight (g) | Morphometry | | | |
|---|---|---|---|---|---|---|
| | | | SL (cm) | TL (cm) | Thickness (cm) | Height (cm) |
| WT1 | M | 0.4682 | 3.1 | 3.9 | 0.5 | 0.7 |
| WT2 | M | 0.4587 | 3.2 | 3.9 | 0.5 | 0.7 |
| WT3 | F | 0.4332 | 3.1 | 3.8 | 0.4 | 0.7 |
| WT4 | F | 0.3759 | 2.7 | 3.3 | 0.3 | 0.7 |
| WT5 | F | 0.4265 | 3.3 | 4 | 0.5 | 0.7 |
| WT6 | F | 0.4529 | 3 | 3.7 | 0.4 | 0.7 |
| WT7 | F | 0.4457 | 3 | 3.6 | 0.3 | 0.7 |
| WT8 | F | 0.5777 | 3.2 | 3.9 | 0.4 | 0.8 |
| WT9 | M | 0.4272 | 3.1 | 3.8 | 0.3 | 0.6 |
| WT10 | M | 0.677 | 3.2 | 4.1 | 0.5 | 0.7 |
| *col22a1*$^{TSPN-/-}$ 1 | M | 0.4620 | 3 | 4 | 0.5 | 0.7 |
| *col22a1*$^{TSPN-/-}$ 2 | M | 0.4202 | 3.3 | 3.8 | 0.5 | 0.7 |
| *col22a1*$^{TSPN-/-}$ 3 | F | 0.3689 | 3.1 | 4.2 | 0.5 | 0.7 |
| *col22a1*$^{TSPN-/-}$ 4 | F | 0.4368 | 3 | 3.7 | 0.5 | 0.7 |
| *col22a1*$^{TSPN-/-}$ 5 | M | 0.3690 | 3 | 3.5 | 0.4 | 0.7 |
| *col22a1*$^{TSPN-/-}$ 6 | M | 0.3698 | 2.8 | 3.8 | 0.4 | 0.7 |
| *col22a1*$^{TSPN-/-}$ 7 | F | 0.7692 | 2.5 | 3.3 | 0.5 | 0.7 |
| *col22a1*$^{TSPN-/-}$ 8 | F | 0.3983 | 3 | 4 | 0.4 | 0.7 |
| *col22a1*$^{TSPN-/-}$ 9 | F | 0.3309 | 3.1 | 3.5 | 0.4 | 0.7 |
| *col22a1*$^{TSPN-/-}$ 10 | F | 0.5845 | 3.1 | 4 | 0.6 | 0.8 |
| *col22a1*$^{TSPN-/-}$ 11 | M | 0.3893 | 2.8 | 3.5 | 0.4 | 0.6 |
| *col22a1*$^{TSPN-/-}$ 12 | F | 0.6041 | 3.4 | 3.7 | 0.4 | 0.7 |
| *col22a1*$^{TSPN-/-}$ 13 | M | 0.4219 | 3 | 3.5 | 0.4 | 0.7 |

**Supplementary videos**

**Video 1:** Swimming capacity of a 2wpf WT larva (*col22a1*$^{TSPN+/+}$). The swimming capacity and postural behavior were recorded at different time intervals using a Zeiss Axiozoom V16 stereoscopic microscope with a digital camera system.

**Video 2:** Swimming capacity of 2wpf of a class 1 mutant larva (*col22a1*$^{TSPN-/-}$). The swimming capacity and postural behavior were recorded at different time intervals using Zeiss Axiozoom V16 stereoscopic microscope with a digital camera system.

**Video 3:** Swimming capacity of 2wpf of a class 2 mutant larva (*col22a1*$^{TSPN-/-}$). The swimming capacity and postural behavior were recorded at different time intervals using a Zeiss Axiozoom V16 stereoscopic microscope with a digital camera system.

**Video 4:** Swimming performance of a 6mpf wildtype sibling (col22a1$^{TSPN+/+}$) fish using a high-speed camera MiroM310. The video was acquired during swimming step protocol experiment at 5BL.s$^{-1}$ water flow and slowed down 10 times vs real speed.

**Video 5:** Swimming performance of a 6mpf class 1 mutant fish (col22a1$^{TSPN-/-}$) using a high-speed camera MiroM310. The video was acquired during swimming step protocol experiment at 5 BL.s-1 water flow and slowed down 10 times *vs* real speed.

**Video 6:** Swimming performance of a 6mpf wildtype sibling (col22a1$^{TSPN+/+}$) fish using a high-speed camera MiroM310. The video was acquired during swimming step protocol experiment at 8BL.s$^{-1}$ water flow and slowed down 10 times vs real speed.

**Video 7:** Swimming performance of a 6mpf class 1 fish (col22a1$^{TSPN-/-)}$ using a high-speed camera MiroM310. The video was acquired during swimming step protocol experiment at 8 BL.s-1 water flow and slowed down 10 times vs real speed.
